# Strong non-ideality effects at low protein concentrations: considerations for elongated proteins

**DOI:** 10.1101/2022.12.22.521448

**Authors:** Alexander E. Yarawsky, Vlad Dinu, Stephen E. Harding, Andrew B. Herr

**Affiliations:** Division of Immunobiology, Cincinnati Children’s Hospital Medical Center, Cincinnati, OH, USA; National Centre for Macromolecular Hydrodynamics (NCMH), University of Nottingham, Sutton Bonington, Loughborough, LE12 5RD UK; Division of Infectious Diseases, Cincinnati Children’s Hospital Medical Center, Cincinnati, OH, USA; Department of Pediatrics, University of Cincinnati College of Medicine, Cincinnati, OH, USA

**Keywords:** Hydrodynamic non-ideality, viscosity, sedimentation velocity, global analysis, hydrodynamic modeling, viscous fingering

## Abstract

A recent investigation was aimed at obtaining structural information on a highly extended protein via SEC-MALS-SAXS. Significantly broadened elution peaks were observed, reminiscent of a phenomenon known as viscous fingering. This phenomenon is usually observed above 50 mg/mL for proteins like bovine serum albumin (BSA). Interestingly, the highly extended protein (Brpt5.5) showed viscous fingering at concentrations lower than 5 mg/mL. The current study explores this and other non-ideal behavior, emphasizing the presence of these effects at relatively lower concentrations for extended proteins. BSA, Brpt5.5, and a truncated form of Brpt5.5 referred to as Brpt1.5 are studied systematically using size-exclusion chromatography (SEC), sedimentation velocity analytical ultracentrifugation (AUC), and viscosity. The viscous fingering effect is quantified using two approaches and is found to correlate well with the intrinsic viscosity of the proteins – Brpt5.5 exhibits the most severe effect and is the most extended protein tested in the study. By AUC, the hydrodynamic non-ideality was measured for each protein via global analysis of a concentration series. Compared to BSA, both Brpt1.5 and Brpt5.5 showed significant non-ideality that could be easily visualized at concentrations at or below 5 mg/mL and 1 mg/mL, respectively. A variety of relationships were examined for their ability to differentiate the proteins by shape using information from AUC and/or viscosity. Furthermore, these relationships were also tested in the context of hydrodynamic modeling. The importance of considering non-ideality when investigating the structure of extended macromolecules is discussed.

## Introduction

There is a long history of interest in the structure of proteins. High-resolution techniques such as X-ray crystallography, NMR, and more recently cryo-EM, are common practice among protein structure labs. Many proteins, however, are not amenable to crystallization, while NMR and cryo-EM require very expensive instrumentation. Additionally, crystallography and cryo-EM approaches require removal of the protein from its native solution environment. This can potentially lead to artifacts related to crystal packing or issues with sample freezing that may provide misleading information regarding tertiary or quaternary structure.

Hydrodynamic approaches may often be neglected, despite their proven usefulness in providing structural insights, and perhaps more importantly, biophysical and mechanistic details. For example, analytical ultracentrifugation (AUC) was developed in the 1920’s by Theodore Svedberg, when he performed sedimentation equilibrium experiments to determine the molecular weight of ovalbumin, hemoglobin, phycocyanin, and phycoerythrin in various buffers (Svedberg and Fåhraeus 1926; Svedberg and Lewis 1928; Svedberg and Nichols 1926; Svedberg and Nichols 1927). There also exists a rich history in the use of viscosity to determine macromolecular shape and flexibility parameters, dating back to Einstein in the early 1900’s (Einstein 1906; Einstein 1911; Harding 1995; Harding 1997).

During a previous structural investigation of a fibril-like protein, very broad and abnormally shaped SEC elution peaks were observed, despite a highly pure and homogeneous sample (Yarawsky et al. 2022). The elution behavior was reminiscent of a phenomenon known as viscous fingering, which is often observed at high concentrations (~50 mg/mL) for proteins like bovine serum albumin (BSA) (Plante et al. 1994). This phenomenon can also be observed when the injected sample is spiked with dextran to increase the viscosity of the sample (Flodin 1961). The root cause of the effect is the fingering that occurs at the interface between a more viscous solution and a less viscous solvent.

To better understand why this protein showed such prominent viscous fingering while at relatively low concentrations (Yarawsky et al. 2022), a systematic analysis of several proteins was performed. The protein construct with which the observations were made previously – Brpt5.5 – is part of the B-repeat superdomain of the biofilm-related accumulation-associated protein (Aap) from *Staphylococcus epidermidis*. The Brpt5.5 construct contains 5 full B-repeats and a C-terminal half-repeat. For comparative purposes, an additional construct containing one and a half B-repeats (Brpt1.5) is also examined here. The Brpt1.5 construct has been extensively studied to show that it exists in solution as an extended monomer (Chaton and Herr 2017; Conrady et al. 2008; Conrady et al. 2013; Shelton et al. 2017). The Brpt5.5 construct has been characterized by extensive hydrodynamic analyses and small-angle X-ray scattering (SAXS), which demonstrated that Brpt5.5 is monomeric and lacked any significant flexibility (Yarawsky and Herr 2020; Yarawsky et al. 2022; Yarawsky et al. 2020). It is worth noting that the biological function of the B-repeat superdomain is to assemble in the presence of Zn^2+^ into amyloid-like fibrils that contribute toward the strength and stability of *S. epidermidis* biofilms. This self-association often requires millimolar concentrations of Zn^2+^ in the context of purified protein in solution and does not assemble as an apo-protein (Chaton and Herr 2017; Shelton et al. 2017; Yarawsky and Herr 2020). In the current study, Brpt1.5 and Brpt5.5 were only studied in the monomeric form. In addition, bovine serum albumin (BSA) was examined as a protein with a globular conformation. This is in stark contrast to the Brpt1.5 and Brpt5.5 proteins, as seen in Figure 1.

**Fig. 1.**
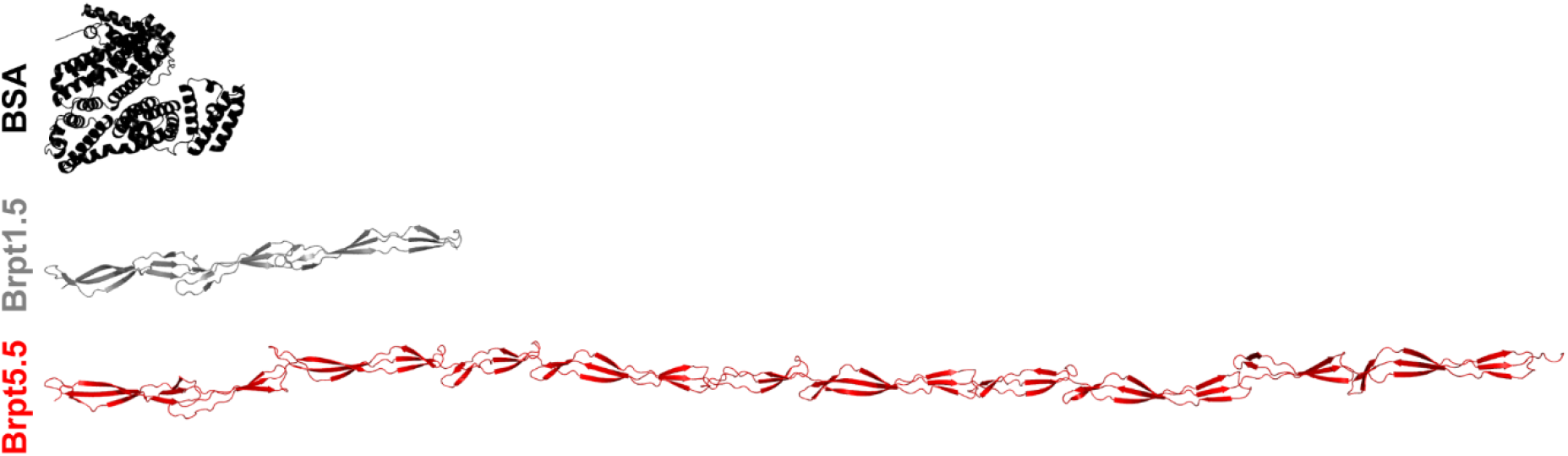
Proteins investigated in the current study. Ribbon models are shown for BSA (PDB: 4f5s), Brpt1.5 (PDB: 4fun), and Brpt5.5 (SASBDB: SASPD43). These images were generated using PyMOL (The PyMOL Molecular Graphics System, version 2.4, Schrödinger, LLC).

This study investigates the impact of shape on hydrodynamic behavior, especially as it relates to size-exclusion chromatography, sedimentation velocity AUC, and viscosity. It demonstrates the clear ability of hydrodynamic techniques to provide structural insights. Additionally, the importance of considering and measuring non-ideality in sedimentation velocity experiments is discussed. Oftentimes, proteins are assumed to behave ideally in “dilute” solutions. However, it is difficult to know at what point it is reasonable to assume that non-ideality is not impacting the data without having collected data at multiple concentrations. Brpt5.5 provides an excellent example of a protein that behaves non-ideally at concentrations below 1 mg/mL – where many investigators may assume an ideal species.

## Results

### Viscous fingering at low concentrations

Each protein was purified and dialyzed into matching 20 mM Tris, 150 mM NaCl (pH 7.4) buffer. Next, the proteins were analyzed by SEC using a Superose 6 (24 mL) column with an injection volume of 250 μL and flow rate of 0.5 mL/min at 4°C. Figure 2 shows the elution profile for (A) BSA, (B) Brpt1.5, and (C) Brpt5.5. While BSA required a 25 mg/mL loading concentration before significant peak broadening occurred, Brpt1.5 and Brpt5.5 showed a similar effect at 10 mg/mL and 4 mg/mL loading concentrations, respectively. To better quantify the severity of the viscous fingering, the height equivalent of a theoretical plate (HETP) and asymmetry factor (*A*_s_) were calculated for each elution (Figure 2D and Figure 2E). These are parameters commonly used to evaluate column performance. Higher values indicate peak broadening and tailing associated with poor column performance. However, the HETP parameter has also been used to quantify viscous fingering effects (Plante et al. 1994). From the results shown in Figure 2, it is very apparent that Brpt5.5 displays viscous fingering effects at much lower concentrations than BSA.

**Fig. 2.**
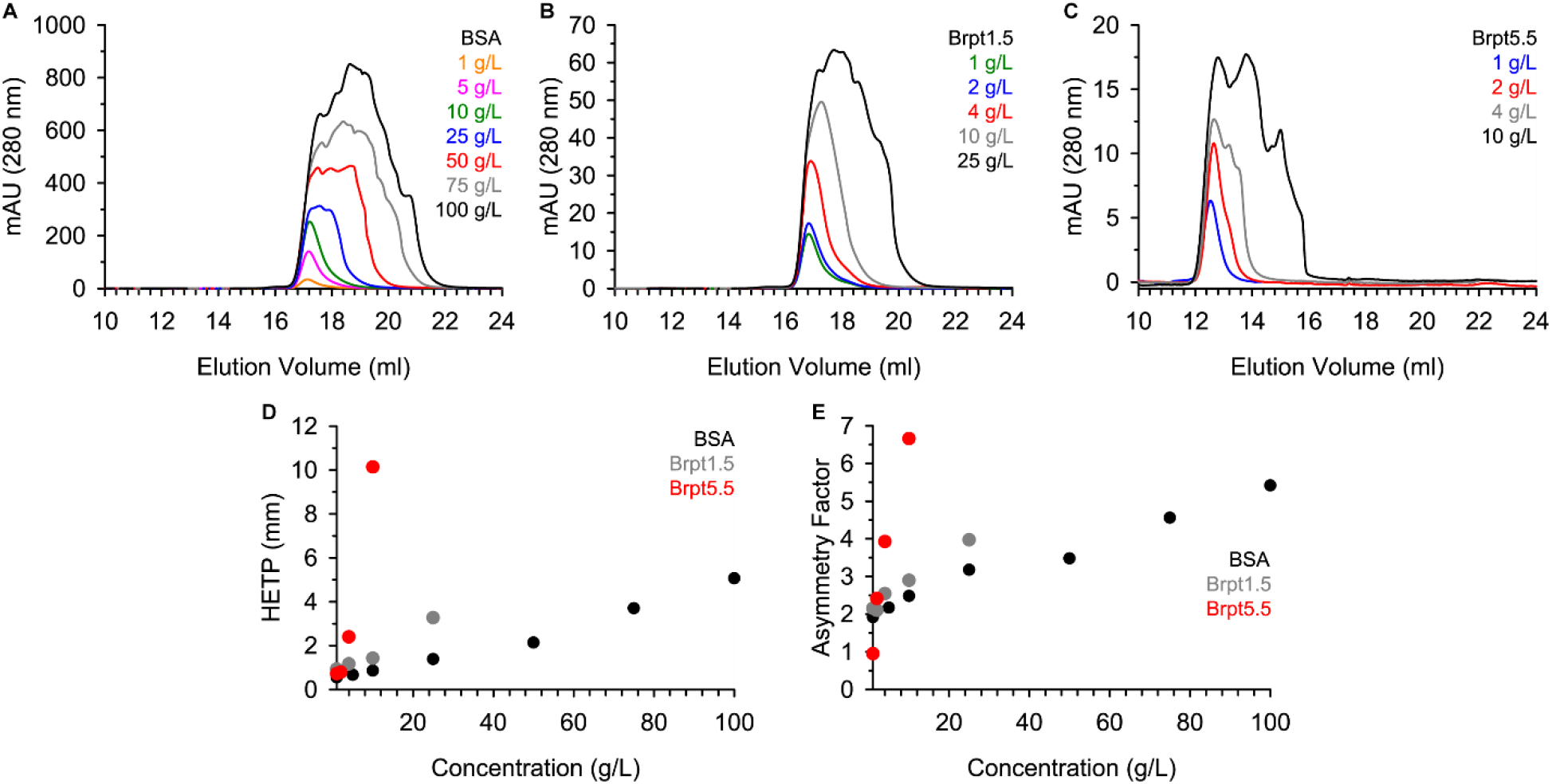
Proteins were analyzed by size-exclusion chromatography. SEC chromatograms are shown for (**A**) BSA, (**B**) Brpt1.5, and (**C**) Brpt5.5 at multiple concentrations. The (**D**) HETP and (**E**) asymmetry factor (*A*_s_) were calculated based on each SEC elution profile as described in the Materials and Methods section. Brpt5.5 SEC data (**C**) were previously published (Yarawsky et al. 2022).

### Intrinsic viscosity correlates with shape

Given that the viscosity of the injected solution is relevant to the severity of viscous fingering, it could be inferred that a solution of Brpt5.5 would have a higher viscosity than a solution of BSA at the same protein concentration. The viscosity of protein solutions may also be directly measured using a capillary viscometer. Figure 3A shows the measured relative viscosity (ηrel) of solutions of BSA, Brpt1.5, and Brpt5.5. A trend was observed that reflected the HETP results very closely, confirming that the viscous fingering effects observed by SEC were in fact due to increased viscosity of Brpt1.5 and Brpt5.5.

**Fig. 3.**
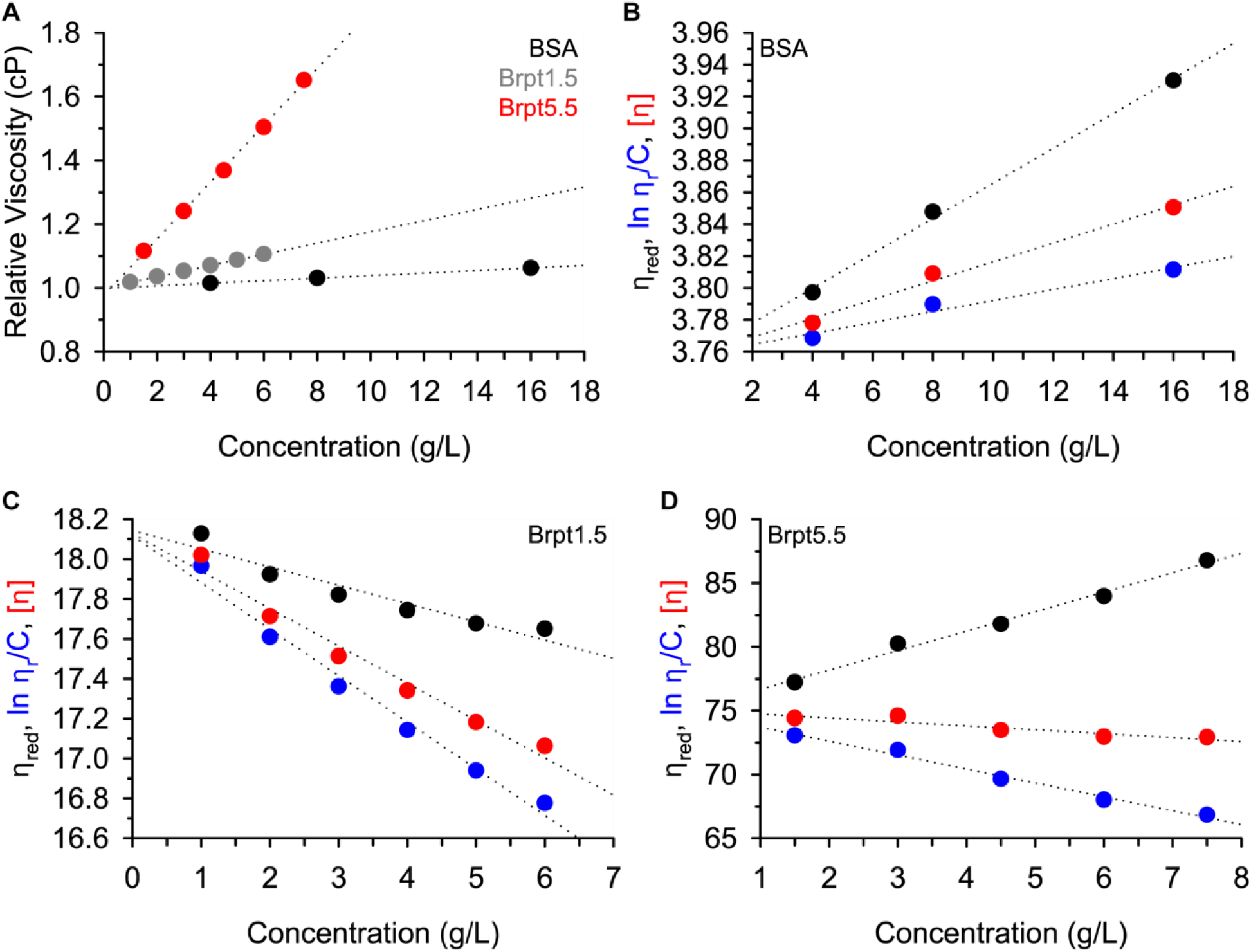
Viscosity was measured for each protein at several concentrations. Panel (**A**) shows the relative viscosity data for all three proteins, with markers colored in accordance with the panel legend (BSA = black, Brpt1.5 = grey, Brpt5.5 = red). The reduced viscosity (black), inherent viscosity (blue), and intrinsic viscosity (red) is plotted for BSA in panel (**B**), Brpt1.5 in panel (**C**), and Brpt5.5 in panel (**D**). In all cases, the dotted lines represent linear fits of each dataset.

The reduced specific viscosity (ηred), inherent viscosity ((ln ηrel)/c), and intrinsic viscosity ([η]) were determined at multiple concentrations, as plotted in Figure 3 for each protein. Extrapolation to zero concentration removes concentration-dependent effects and provides the fundamental macromolecular information. The intrinsic viscosity is particularly useful in its relationship to macromolecular conformation, flexibility, and hydration. Macromolecules that exhibit high axial ratios also show high intrinsic viscosities (Creeth and Knight 1965; Harding 1997). The intrinsic viscosities measured for Brpt1.5 and Brpt5.5 are indeed much greater than BSA, correlating well with the more extended shapes of Brpt1.5 and Brpt5.5.

### Hydrodynamic non-ideality measured by sedimentation velocity AUC

High intrinsic viscosity is correlated with an increase in hydrodynamic non-ideality term, *ks,* for extended macromolecules (Creeth and Knight 1965; Harding 1997). The hydrodynamic non-ideality term is related to the backflow of solvent displaced by the macromolecule as it sediments through solution in a closed system, and it is a concentration-dependent phenomenon (Correia and Stafford 2015). Sedimentation velocity experiments were performed across a concentration range to determine the hydrodynamic non-ideality of each protein.

Figure 4 shows g(*s*\*) distributions and WDA (wide distribution analysis) distributions obtained from sedimentation velocity experiments. The g(*s*\*) analysis requires a short time span of data to be used, while the WDA uses all scans collected during the experiment. Both are based on the time-derivative of the concentration profile and are considered model-independent analyses of the data (Philo 2000; Sherwood and Stafford 2016; Stafford 1992). For a case such as this, where non-ideality is of interest, a model-independent approach is preferred over the c(*s*) analysis that assumes ideal, noninteracting species (Schuck 2000). In both the g(*s*\*) and WDA distribution, the effect of hydrodynamic non-ideality is visually apparent. In accordance with Equation 1 and Equation 2, an increase in the *k*_s_*c* term results in a slowing and sharpening of the sedimentation boundary (Correia et al. 2020).

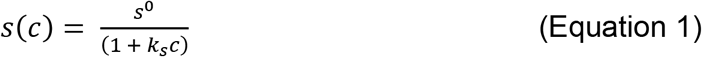

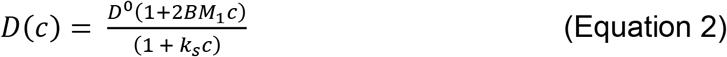

**Fig. 4.**
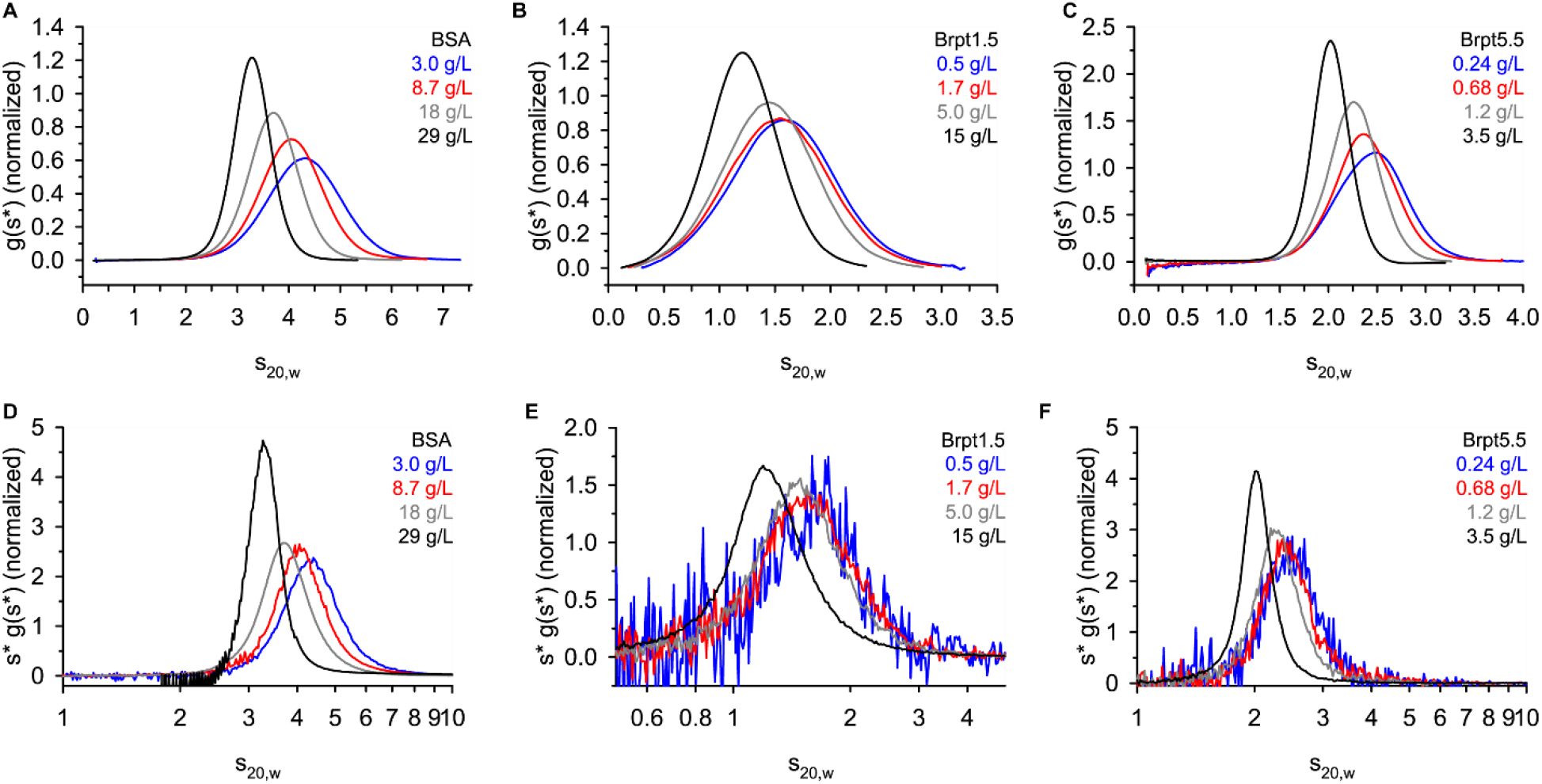
Sedimentation velocity AUC data were analyzed by DCDT+ to obtain a g(*s*\*) distribution (panel **A**-**C**) or by SEDANAL WDA (panel **D**-**F**). Each panel lists the target loading concentration.

Once again, the concentration of Brpt1.5 and Brpt5.5 required to observe a similar effect to BSA is much lower. For example, a significant shift in the g(*s*\*) distribution is observed for BSA at 3 – 8.7 mg/mL, while just 0.68 mg/mL of Brpt5.5 showed a significant shift.

Two basic approaches exist to quantify the *k*_s_ value. The first approach directly fits the relationship between the sedimentation coefficient (*s*_20,w_) and the loading concentration (Correia et al. 2016; Correia et al. 2020; Winzor et al. 2021). The 1/*s*_20,w_ vs concentration plots are shown in Figure 5, along with a standardized plot to show the relative impact of hydrodynamic non-ideality on the sedimentation coefficient. The extreme effect of hydrodynamic non-ideality on the sedimentation of Brpt5.5 is easily visualized in Figure 5D.

**Fig. 5.**
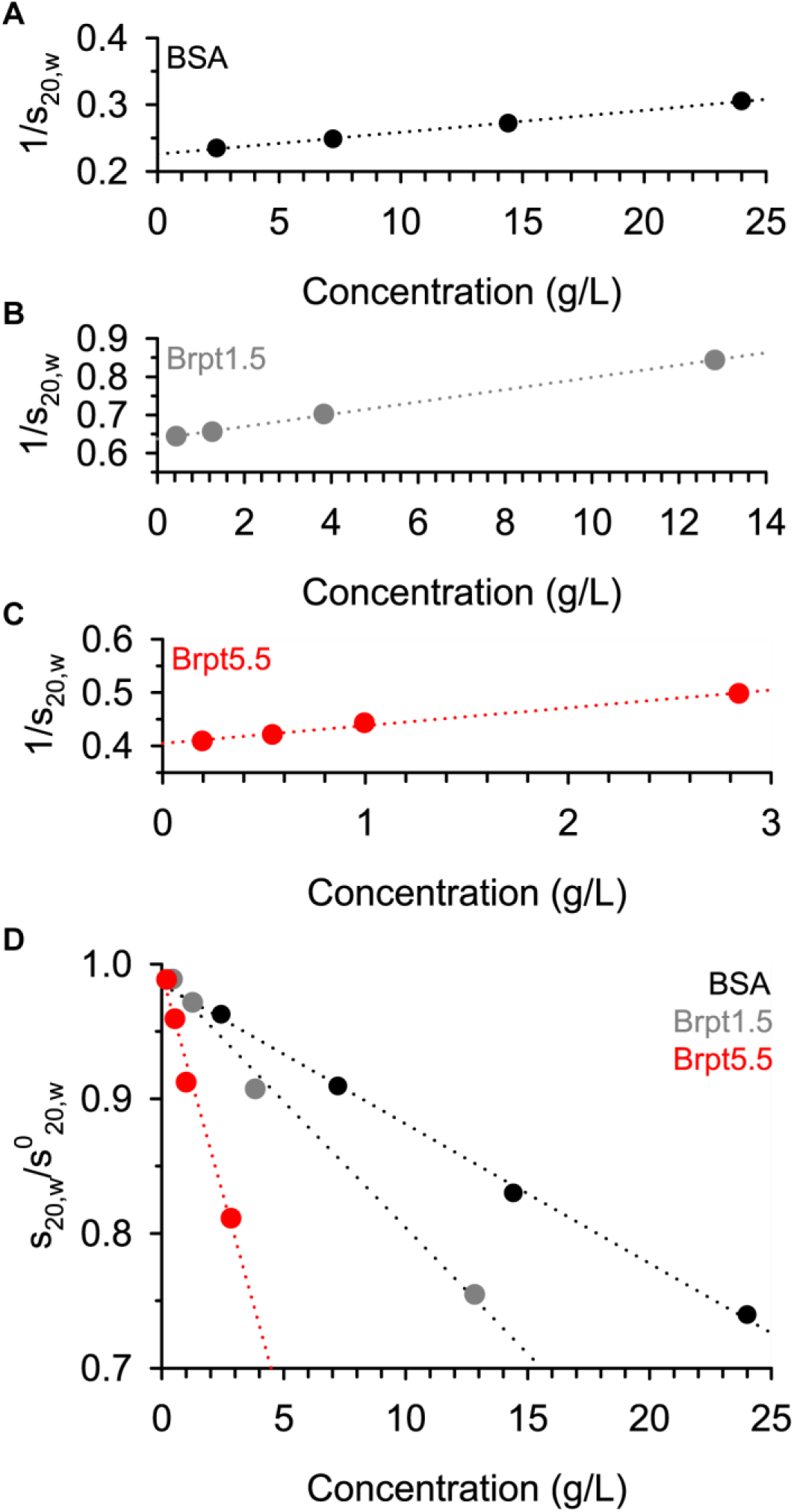
Determination of the concentration-dependence of the sedimentation coefficient was performed within SEDNTERP. Sedimentation coefficients used were those reported by integration in DCDT+ after converting to *s*_20,w_.

A more rigorous approach is to globally fit the data using direct boundary fitting in SEDANAL (Correia et al. 2020; Sherwood and Stafford 2016). Because sedimentation and diffusion both impact the sedimentation boundaries, a model containing both the hydrodynamic non-ideality (*k*_s_) and thermodynamic non-ideality (B*M*_1_) terms should be fitted. The thermodynamic non-ideality term is expected to be on the same order of magnitude as *k*_s_ but is less well-determined by velocity experiments – especially when experiments are run at a high speed to minimize the impact of diffusion with respect to sedimentation. The results of SEDANAL fitting to a non-ideal model are listed in Table 1 and are comparable to the result from linear fitting of the *s*_20,w_ vs concentration data. Figure S1 – Figure S3 shows the quality of fit for each global analysis. Characterization of the concentration-dependence of the sedimentation coefficient also yields the *S*^0^_20,w_ value. The *k*_s_ values are again increasing with protein shape (BSA < Brpt1.5 < Brpt5.5). The especially high *k*_s_ value of Brpt5.5 helps explain why the g(*s*\*) distribution shows significant slowing and sharpening even below 1 mg/mL (Figure 4).

**Table 1.** Analysis of non-ideal sedimentation

|  | SEDNTERP |  | SEDANAL |  |  |
| --- | --- | --- | --- | --- | --- |
| | $s_{20,w}^0$ | $k_s$ (mL/g) | $s_{20,w}^0$ | $k_s$ (mL/g) | $BM_1$ (mL/g) |
| BSA | 4.43<br>(4.41 – 4.45) | 14.5<br>(14.1 – 14.9) | 4.29<br>(4.28 – 4.30) | 14.7<br>(14.6 – 14.9) | 3.0<br>(2.5 – 3.5) |
| Brpt1.5 | 1.57<br>(1.57 – 1.58) | 25.5<br>(25.0 – 26.0) | 1.54<br>(1.54 – 1.54) | 23.5<br>(23.5 – 23.6) | 12.5<br>(12.4 – 12.7) |
| Brpt5.5 | 2.48<br>(2.46 – 2.50) | 84.0<br>(77.8 – 90.2) | 2.30<br>(2.30 – 2.31) | 68.0<br>(67.5 – 68.5) | 4.4<br>(0.5 – 8.6) |

SEDNTERP error limits are $\pm 1$ standard deviation.
SEDANAL error limits are an estimate of the 95% confidence interval using F-statistics.

### Determination of shape from AUC

A parameter of interest for biophysicists and structural biologists alike is the shape of the macromolecule. In the absence of high-resolution structural data, sedimentation velocity AUC experiments can provide very reliable indications of the macromolecule’s global conformation in solution. The Svedberg Equation and Stokes-Einstein Equation relate the sedimentation coefficient (*s*), the diffusion coefficient (*D*), the buoyant mass (*M*_b_), and the frictional coefficient (*f*) (Chaton and Herr 2015; Correia and Stafford 2015; Rocco and Byron 2015b). Rather than reporting the frictional coefficient, the more useful parameter is the frictional ratio (*f*/*f*_0_). This is the ratio of the frictional coefficient of the solute to the frictional coefficient of an ideal sphere of the same volume and offers a shape description that is independent of the macromolecule’s size. Accurate determination of the *f*/*f*_0_ relies on accurate fitting of the concentration-dependence of *s* and *D*. Where non-ideality is present, this means also fitting both the hydrodynamic and thermodynamic non-ideality terms.

A common approach to determining *f*/*f*_0_ is to utilize the c(*s*) analysis implemented in SEDFIT. However, considering the major concentration-dependence of both *s* and *D* due to non-ideality, global fitting to a non-ideal model is required. For illustrative purposes, Table S1 lists the *s*, *M,* and *f*/*f*_0_ for each individual dataset analyzed by c(*s*). The higher concentration datasets could be interpreted as highly extended dimer or trimers. A similar discrepancy is observed when fitting g(*s*\*) distributions in DCDT+. This is simply due to the impact of *k*_s_*c* (and B*M*_1_c) on *s* and *D*. It should be noted that the quality of these fits to ideal models is quite poor, so careful analysis by the user should avoid such erroneous results from being reported.

Direct boundary fitting performed in SEDANAL to the non-ideal model used previously was therefore used to determine the *f/f*_0_ (at infinite dilution). An alternative approach would be to use the fitted *s*^0^_20,w_ and *M* to calculate *f*/*f*_0_. The SEDANAL-fitted *f*/*f*_0_ was incorporated into Table 2 with error estimates for each protein. Using X-ray crystallography structures of BSA and Brpt1.5, and a SAXS-validated structure of Brpt5.5, a comparison of the experimentally determined *f*/*f*_0_ and theoretical *f*/*f*_0_ from hydrodynamic modeling could be compared. Good agreement with hydrodynamic modeling via HullRad (Fleming and Fleming 2018) (Table 3) indicates that AUC can be effectively used to estimate the shape of highly non-ideal macromolecules.

**Table 2.** Shape parameters determined in the study

|  |  |  |  |  |  |  |  |
| --- | --- | --- | --- | --- | --- | --- | --- |
| | $M$ (Da) | $\bar{v}$ (mL/g) | | $[\eta]$ | $\nu$ | $s^0_{20,w}$ | $f/f_0$ |
| BSA | 66,398.00 | 0.7325 |  | 3.8 | 5.2 | 4.29 | 1.3 |
| Brpt1.5 | 22,217.26 | 0.7292 |  | 18.1 | 24.8 | 1.54 | 1.8 |
| Brpt5.5 | 78,030.99 | 0.7280 |  | 75.1 | 103.2 | 2.30 | 2.7 |
| | Hydration<br>(g/g) | P | $a/b$ (P) | $\beta$ (x10<sup>6</sup>) | $a/b$ ($\beta$) | $k_s$ | $k_s/[\eta]$ |
| BSA | 0.441 | 1.1 | 3.0 | 1.96 | 1.0 | 14.7 | 3.9 |
| Brpt1.5 | 0.516 | 1.5 | 9.3 | 2.45 | 11.7 | 23.5 | 1.3 |
| Brpt5.5 | 0.526 | 2.3 | 26.9 | 2.53 | 15.1 | 68.0 | 0.9 |

$M$ is the theoretical molar mass based on amino acid composition.
$\bar{v}$ and hydration are based on amino acid composition and reported by SEDNTERP.
For experimental values, $a/b$ (P) was based on the Perrin function, while $a/b$ ( $\beta$ ) was based on the Scheraga-Mandelkern $\beta$ function.
$s_{20,w}^0$ and $k_s$ values reported by SEDANAL global fitting to a non-ideal monomer model.

**Table 3.** Comparison of experimental results to hydrodynamic modeling

|  | Experimental |  |  |  | HullRad |  |  | AtoB/ZENO |  |  |  |
| --- | --- | --- | --- | --- | --- | --- | --- | --- | --- | --- | --- |
| | $s^0_{20,w}$ | $f/f_0$ | $a/b$<br>(P) | $a/b$<br>($\beta$) | $s^0_{20,w}$ | $f/f_0$ | $a/b$ | $s^0_{20,w}$ | $f/f_0$ | $a/b$ | $[\eta]$ |
| BSA | 4.29 | 1.3 | 3.0 | 1.0 | 4.39 | 1.3 | 1.5 | 4.59 | 1.3 | 1.4 | 3.9 |
| Brpt1.5 | 1.54 | 1.8 | 9.3 | 11.7 | 1.64 | 1.7 | 7.3 | 1.71 | 1.7 | 7.0 | 13.7 |
| Brpt5.5 | 2.30 | 2.7 | 26.9 | 15.1 | 2.58 | 2.6 | 16.0 | 2.47 | 2.7 | 15.6 | 92.1 |

Models shown in Figure 1 were used for hydrodynamic modeling.
For experimental values, $a/b$ (P) was based on the Perrin function, while $a/b$ ( $\beta$ ) was based on the Scheraga-Mandelkern $\beta$ function.
For AtoB/ZENO calculations, $\bar{v}$ was based on amino acid composition (SEDNTERP).

### Evaluation of shape parameters using viscosity and AUC data

Many studies have been performed utilizing *either* hydrodynamic data from AUC *or* viscosity data have been utilized to gain structural insights into macromolecules. Because both sets of data have been collected and structures have already been determined for the proteins of interest, the current study provides an excellent opportunity to evaluate various relationships often used in the field. Creeth and Knight published a seminal review focused on estimation of shape from sedimentation and viscosity data (Creeth and Knight 1965). Since then, additional work has been focused on further developing these approaches and has been summarized by Harding (Harding 1995).

The ratio of intrinsic viscosity ([η]) over partial specific volume (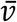 or “v-bar”) known as the viscosity increment (*v*) is expected to yield a ratio of 2.5 for spheres but increases with asymmetry. The *f/f*_0_ has already been discussed above, where a value of 1.0 would indicate an anhydrous, ideal hard sphere. Globular proteins often are expected to exhibit a *f*/*f*_0_ ~1.2-1.3. More specific shape information can be obtained from the Perrin function (P). This parameter distinguishes between contributions of molecular shape and hydration (requiring hydration information), and it can be used to define the axial ratio *(a/b)* of a prolate or oblate ellipsoid (Harding 1995) using the ELLIPS1 software (Harding et al. 1997). Equations describing both the viscosity increment and *f*/*f*_0_ can be used to produce the Scheraga-Mandelkern *β* function. The result is independent of hydration, but requires very accurate experimental determination of *s*, [η], *M,* and 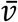, and is often considered to be rather insensitive to shape (Creeth and Knight 1965; Harding 1995).

Analyses of *k*_s_ and [η] by Wales and van Holde indicate that *k*_s_/[η] provides a shape parameter that is sensitive to global conformation and does not require any assumptions about hydration. A ratio of ~1.6 is expected for globular proteins, while asymmetry pushes the ratio lower. The Wales-van Holde ratio is a highly attractive parameter to describe shape, given that both *k*_s_ and [η] can be readily measured.

With high-resolution structures or SAXS-based models available for the three proteins, it is possible to perform hydrodynamic bead modeling (Bujacz 2012; Conrady et al. 2013; Yarawsky et al. 2022). This allows for comparison of predicted vs experimental *s*^0^_20,w_, *f*/*f*_0_, and *a/b* values from HullRad (Fleming and Fleming 2018). An additional approach was taken using the AtoB bead modeling approach with ZENO hydrodynamic calculations, as implemented in US-SOMO (Brookes et al. 2010; Byron 1997; Juba et al. 2017; Kang et al. 2004; Rocco and Byron 2015a). Overall, the hydrodynamic predictions are in reasonable agreement with the experimental parameters. The major discrepancy is in the *a/b* ratio derived from the Perrin (P) function, which severely overestimated the degree of extension in the molecules in reference to the estimate by the Scheraga-Mandelkern *β* function and hydrodynamic modeling. Hydrodynamic predictions tend to be especially useful when evaluating the plausibility of several different conformational models, rather than describing the exact shape and dimensions of a given molecule in solution (Brautigam et al. 2020; Marx et al. 2020; Monsen et al. 2021; Rocco and Byron 2015b; Yarawsky et al. 2022). Despite minor inconsistencies in the hydrodynamic predictions and experimental data, it is very clear that there is increasing extension from BSA to Brpt1.5 and Brpt5.5.

## Discussion

The goal of this study was to examine the interplay between viscous fingering in chromatography and non-ideal sedimentation. Viscosity measurements demonstrated that the aberrant elution profiles observed, especially for Brpt5.5 at concentrations below 5 mg/mL, were indeed due to high solution viscosity. The increased viscosity of Brpt5.5 also correlated with strong hydrodynamic non-ideality measured by sedimentation velocity AUC, which presents itself via a slowing and sharpening of the g(*s*\*) distribution. An advantage of including Brpt1.5 in this study was to demonstrate the effect of increased axial ratio, without significantly changing the secondary structure or amino acid composition of the protein. Similar comparisons have been performed with fractionated collagen (Nishihara and Doty 1958), however, the present study is able to ensure homogeneity in the samples. In agreement with the collagen data, the longer molecules exhibited higher intrinsic viscosity and hydrodynamic non-ideality (Creeth and Knight 1965). BSA acted as a globular reference and showed results more consistent with a spherical particle. Indeed, the direct ratio of *k*_s_/[η] provided a simple indication of extended shape in Brpt1.5 and Brpt5.5, compared to BSA.

Not only does this study demonstrate the effects of shape on non-ideality and viscosity, but it also provides an opportunity to evaluate the usefulness of certain shape parameters and relationships that have been previously proposed and used within the field. A common result presented with sedimentation velocity data is the *f*/*f*_0_. A value of ~1.3 is expected for globular proteins, while higher values are expected with higher asymmetry. A more explicit shape parameter for ellipsoids is the *a/b* (axial) ratio. Table 2 lists the *a/b* ratio derived from the Perrin function and the Scheraga-Mandelkern β function. In comparison with hydrodynamic modeling based on the protein structures, the *a/b* ratio from the hydration-independent β function is in better agreement than that derived from the Perrin function. Though the β function itself has been considered relatively insensitive to shape changes (for example, 2.45 x 10^6^ vs 2.53 x 10^6^ for Brpt1.5 and Brpt5.5) the *a/b* ratio derived can still prove to be quite useful. Nevertheless, both derivations of the *a/b* ratio indicate much greater extension in the Brpt1.5 and Brpt5.5 proteins than in BSA.

While hydrodynamic approaches are unable to provide high-resolution structural information like X-ray crystallography or cryo-EM, they do provide relatively easy access to lower-resolution information. In many cases, it may be sufficient to simply exclude given conformations or models for a protein of interest. To demonstrate this concept, hydrodynamic modeling was performed using HullRad and AtoB/ZENO. Both approaches yielded *a/b* ratios similar to those derived from the β function and *f*/*f*_0_ values similar to those experimentally determined. Nonetheless, it is evident how hydrodynamic modeling may be able to provide sufficient information to exclude certain structural models. With the recent advent of new computational approaches for structure prediction such as AlphaFold (Jumper et al. 2021) or Rosetta (Du et al. 2021), it may become more important than ever to understand the capabilities of hydrodynamic approaches like AUC and viscosity measurements, as these provide a relatively simple and effective platform to validate predicted structures.

## Materials and Methods

### Protein Purification

Bovine serum albumin Fraction V (BP1605-100) was resuspended in 20 mM Tris pH 7.4, 150 mM NaCl and purified via Superdex 200 16/600 pg (Cytiva) to yield pure monomeric BSA. Brpt1.5 was composed of amino acids 2017-2223 from Aap (accumulation-associated protein) from *S. epidermidis* strain RP62A. The protein was expressed as a fusion with a His-MBP tag, which was cleaved via TEV protease as previously described (Conrady et al. 2008). Brpt1.5 was additionally purified by anion exchange using a HiTrap ANX FF 5 mL column (Cytiva). The running buffer was 20 mM Tris pH 7.4, 50 mM NaCl, and an elution gradient was produced using 20 mM Tris pH 7.4, 1 M NaCl. Brpt5.5 was composed of amino acids 1505-2223 from Aap from RP62A and contained an N-terminal His-MBP tag and a C-terminal Strep-II tag. A TEV protease cleavage site was present downstream of the N-terminal tags and upstream of the C-terminal tag (Yarawsky et al. 2022; Yarawsky et al. 2020). The purification of Brpt5.5 was performed as previously described (Yarawsky et al. 2022).

### Size Exclusion Chromatography

All SEC data presented were collected using a Superose 6 Increase 10/300 24 mL column (Cytiva) connected to an ÄKTA FPLC system (Cytiva) at 4°C. The column was equilibrated in running buffer (20 mM Tris pH 7.4, 150 mM NaCl). The flow rate was set to 0.5 mL/min and a 250 μL sample was injected. Samples were diluted from a concentrated stock that had been dialyzed overnight into running buffer at 4°C. SEC data were exported from UNICORN and the HETP and *A*_s_ values determined manually using Excel and GUSSI (Brautigam 2015). The HETP value was calculated according to Equation 3, where *L* is the column length (300 mm) and *N* is the number of theoretical plates, calculated according to the Exponentially Modified Gaussian Model shown in Equation 4 (Jeansonne and Foley 1991).

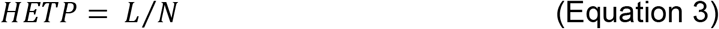

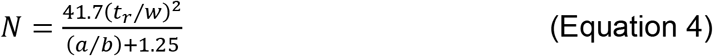

The calculation of *N* used was chosen over other methods to best accommodate asymmetric peaks observed in the presence of viscous fingering. The retention time (*t*_r_) was taken as the weight-average time of the peak of the least concentrated sample. The peak width (*w*) was the width at 10% of the peak height, with *a* and *b* being the distance from tr and the leading (*a*) or trailing edge (*b*) of the peak at 10% max height. The ratio of *b/a* determined the asymmetry (*A*_s_) value. SEC protein standards (BioRad; #1511901) were used to ensure Superose 6 column performance was adequate (Figure S4). A 100 μL aliquot was injected, while all other parameters were kept equivalent to other the experiments.

### Analytical Ultracentrifugation

A Beckman Coulter ProteomeLab XL-I was used to collect interference data at multiple concentrations. Meniscus-matching centerpieces (SpinAnalytical) were used. Data was collected at 48,000 rpm on 4 cells per experiment in an 8-hole rotor (An 50Ti). A radial calibration was performed prior to each experiment. Data collection was monitored using SEDVIEW (Hayes and Stafford 2010). Experiments were carried out until there was no further sedimentation occurring by visual evaluation. To ensure the highest scan frequency and best possible data for g(*s*\*) and SEDANAL fitting, no time delay between scans was used. To ensure scans be collected throughout the entire sedimentation process, the Equilibrium option was chosen rather than the Velocity option within ProteomeLab. A Method schedule of 100 steps of 99 scans, totaling 9900 scans per cell, instead of the standard 999 scans maximum would be collected. The experiment was stopped manually once sufficient data were collected. Analyses were performed in DCDT+ (Philo 2000; Stafford 1992), SEDANAL (Sherwood and Stafford 2016; Stafford and Braswell 2004; Stafford and Sherwood 2004), and SEDFIT (Schuck 2000). Linear fitting of 1/*s*_20,w_ vs concentration to determine *k*_s_ and *s*^0^_20,w_ was performed in SEDNTERP v3 (Laue et al. 1992)[cite J. Philo SEDNTERP paper, current issue]. SEDNTERP v3 was also used to estimate buffer density and viscosity, as well as the 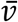, mass, and extinction coefficients for each protein. Within DCDT+, the g(*s*\*) distributions were converted to *s*_20,w_, and the concentration (fringes) and weight-average *s* value from the time of analysis 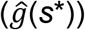 were used for *k*_s_ and *s*^0^_20,w_ determination. The resting rotor temperature of the AUC was determined using a NIST-calibrated DS1922L iButton (iButtonLink) following a procedure previously described (Ghirlando et al. 2014). The measured temperature was incorporated into SEDNTERP for more accurate estimates of the buffer and solute parameters. Error analysis of SEDANAL fits was performed using F-Statistics at a 95% confidence level. The following parameters were allowed to fit: loading concentration, *s*, *k*_s_, B*M*_1_, and the molar mass or frictional ratio.

### Viscosity Measurements

A semi-automated U-tube Ostwald capillary viscometer (Schott Geräte, Hofheim, Germany) was used to measure buffer (*t*_0_) and sample solution (*t*_s_) flow times. Aliquots of 2 mL were prepared at multiple concentrations from dilution of a stock solution, and the relative viscosity (η_r_) was calculated according to Equation 5:

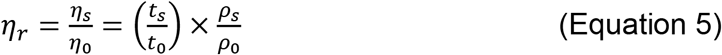

which includes the buffer (ρ_0_) and solution (ρ_s_) density and the buffer (η_0_) and solution (η_s_) viscosity. The buffer and sample solution density values were assumed to be equivalent. All viscosity experiments were performed at 20°C using samples dialyzed into the same buffer (20 mM Tris pH 7.4, 150 mM NaCl).

The reduced viscosity (η_red_) was determined according to the Huggins Equation (Huggins 1942) by extrapolation to zero concentration (*c*).

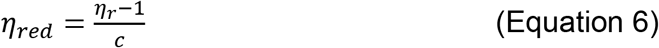

The Kraemer Equation (Kraemer 1938) was used to describe the inherent viscosity (η_inh_) at each concentration:

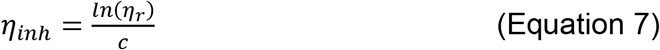

Lastly, the Solomon-Ciuta Equation (Solomon and Ciută 1962) was used to determine the intrinsic viscosity ([η]), where η_sp_ is the reduced specific viscosity (η_rel_ – 1):

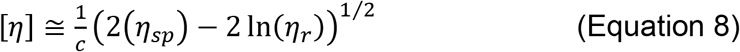

## Author contributions

A.E.Y. conceived the project, designed experiments, performed experiments, analyzed data, and wrote the manuscript

V.D. designed experiments, performed experiments, and analyzed data

S.E.H. guided analysis and reviewed the manuscript

A.B.H. designed experiments and provided funding

All authors reviewed the manuscript and contributed significantly to the final product.

