## Supplemental Information for "Strong non-ideality effects at low protein concentrations: considerations for elongated proteins"

**Fig S1-S4**

**Table S1**

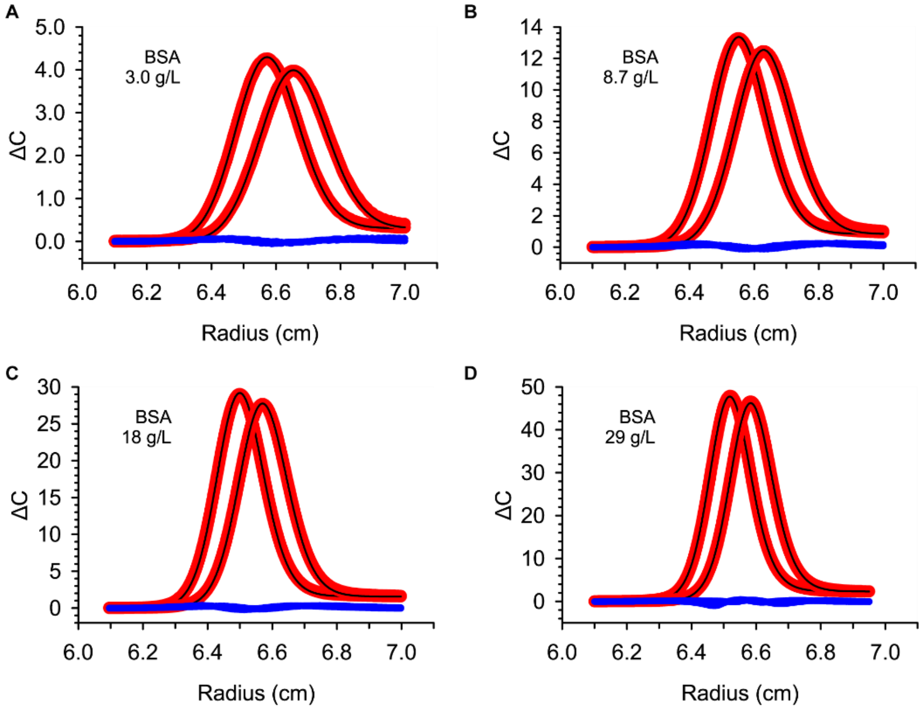


**Fig. S1** SEDANAL global fitting of BSA sedimentation velocity data. A single species, non-ideal model incorporating both $k_s$ and $BM_1$ was used. Concentrations listed in the panels are the target loading concentrations.

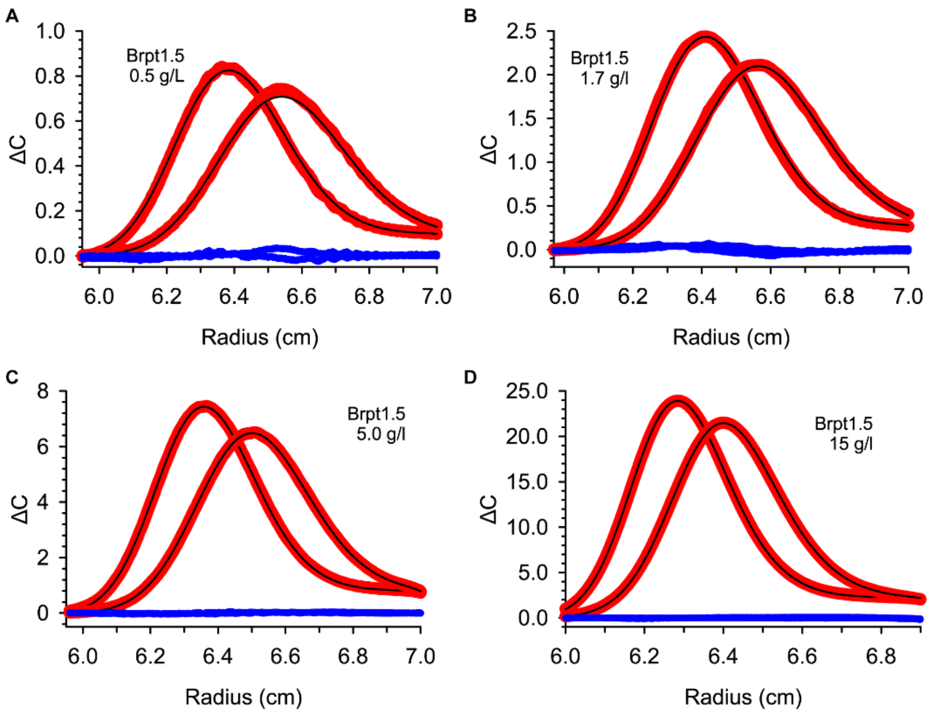


**Fig. S2** SEDANAL global fitting of Brpt1.5 sedimentation velocity data. A single species, non-ideal model incorporating both $k_s$ and $BM_1$ was used. Concentrations listed in the panels are the target loading concentrations.

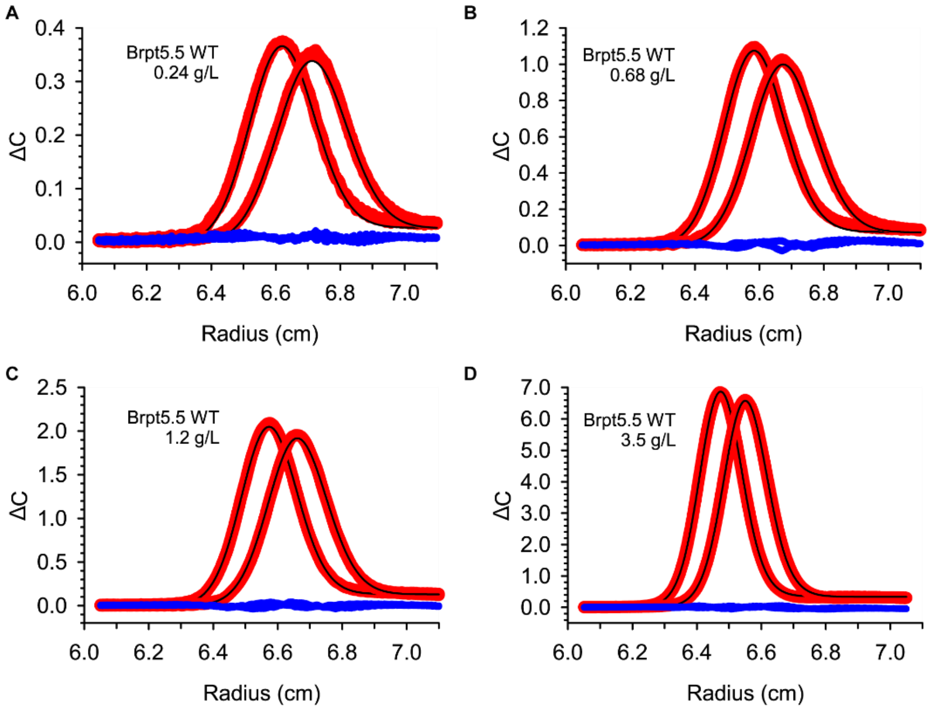


**Fig. S3** SEDANAL global fitting of Brpt5.5 sedimentation velocity data. A single species, non-ideal model incorporating both $k_s$ and $BM_1$ was used. Concentrations listed in the panels are the target loading concentrations.

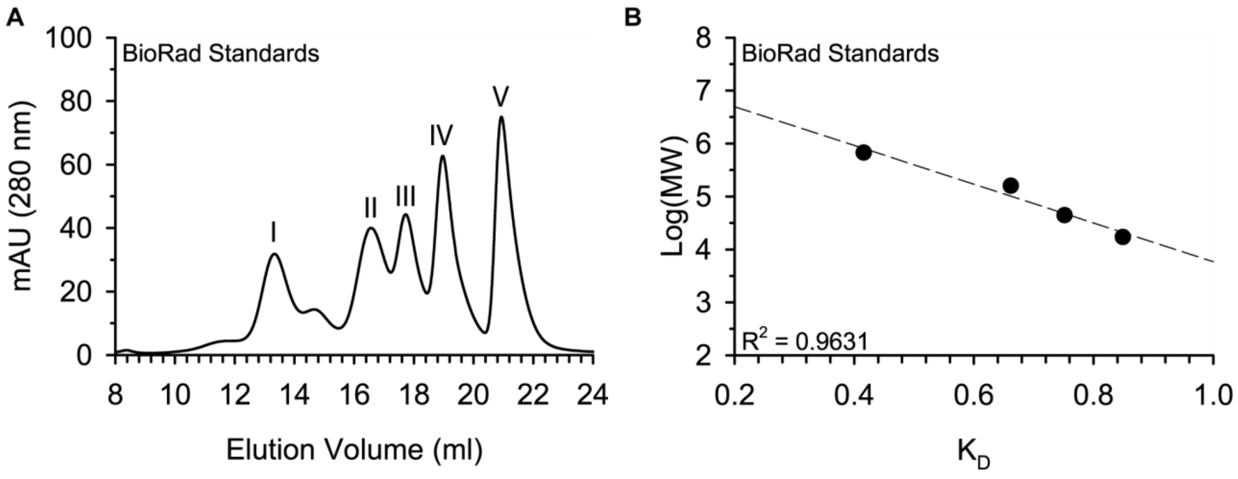


**Fig. S4** SEC chromatogram of protein standards (**A**). The protein standards included thyroglobulin (I), gamma-globulin (II), ovalbumin (III), myoglobulin (IV), and vitamin B12 (V). Panel (**B**) shows the relationship between $K_D$ (retention factor) and the Log(*MW*) of the standards.

**Table S1** Results from local fitting of Brpt5.5 sedimentation velocity data.

| | SEDFIT c(*s*) | | DCDT+ g(*s**) |
|---|---|---|---|
| Conc (g/L) | *M* (kDa) | $f/f_0$ | *M* (kDa) |
| 3.56 | 213 | 6.4 | 227 |
| 1.23 | 114 | 3.8 | 123 |
| 0.68 | 97 | 3.2 | 101 |
| 0.24 | 85 | 2.8 | 87 |

Sequence-based mass of Brpt5.5 is 78.0 kDa.
